# DeepG4 : A deep learning approach to predict active G-quadruplexes from DNA

**DOI:** 10.1101/2020.07.22.215699

**Authors:** Vincent Rocher, Matthieu Genais, Elissar Nassereddine, Raphael Mourad

## Abstract

DNA is a complex molecule carrying the instructions an organism needs to develop, live and reproduce. In 1953, Watson and Crick discovered that DNA is composed of two chains forming a double-helix. Later on, other structures of DNA were discovered and shown to play important roles in the cell, in particular G-quadruplex (G4). Following genome sequencing, several bioinformatic algorithms were developed to map G4s in vitro based on a canonical sequence motif, G-richness and G-skewness or alternatively sequence features including k-mers, and more recently machine/deep learning. Here, we propose a novel convolutional neural network (DeepG4) to map active G4s (forming both in vitro and in vivo). DeepG4 is very accurate to predict active G4s, while most state-of-the-art algorithms fail. Moreover, DeepG4 identifies key DNA motifs that are predictive of G4 activity. We found that active G4 motifs do not follow a very flexible sequence pattern as current algorithms seek for. Instead, active G4s are determined by numerous specific motifs. Moreover, among those motifs, we identified known transcription factors (TFs) which could play important roles in G4 activity by contributing either directly to G4 structures themselves or indirectly by participating in G4 formation in the vicinity. Moreover, we showed that specific TFs might explain G4 activity depending on cell type. Lastly, variant analysis suggests that SNPs altering predicted G4 activity could affect transcription and chromatin, *e.g*. gene expression, H3K4me3 mark and DNA methylation. Thus, DeepG4 paves the way for future studies assessing the impact of known disease-associated variants on DNA secondary structure by providing a mechanistic interpretation of SNP impact on transcription and chromatin.

Availability: https://github.com/morphos30/DeepG4.

**Author summary:** DNA is a molecule carrying genetic information and found in all living cells. In 1953, Watson and Crick found that DNA has a double helix structure. However, other DNA structures were later identified, and most notably, G-quadruplex (G4). In 2000, the Human Genome Project revealed the widespread presence of G4s in the genome using algorithms. To date, all G4 mapping algorithms were developed to map G4s on naked DNA, without knowing if they could be formed in the cell. Here, we designed a novel artificial intelligence algorithm that could map G4s active in the cell from the DNA sequence. We showed its better accuracy compared to existing algorithms. Moreover, we identified key transcriptional factor motifs that could explain G4 activity depending on cell type. Lastly, we demonstrated the existence of mutations that could alter G4 activity and therefore impact molecular processes, such as transcription, in the cell. Such results could provide a novel mechanistic interpretation of known disease-associated mutations.

## 1 Introduction

Deoxyribonucleic acid (DNA) is a complex molecule carrying genetic instructions for the development, functioning, growth and reproduction of all known living beings and numerous viruses. In 1953, Watson and Crick discovered that DNA is composed of two chains forming a double-helix [44]. However, other structures of DNA were discovered later and shown to play important roles in the cell. Among those structures, G-quadruplex (G4) was discovered in the late 80’s [38]. G4 sequence contains four continuous stretches of guanines [10]. Four guanines can be held together by Hoogsteen hydrogen bonding to form a square planar structure called a guanine tetrad (G-quartets). Two or more G-quartets can stack to form a G4 [10]. The quadruplex structure is further stabilized by the presence of a cation, especially potassium, which sits in a central channel between each pair of tetrads [6]. G4 can be formed of DNA [40] or RNA [17]. Moreover, non-G-quadruplexes also exist [34].

G4s were found enriched in gene promoters, DNA replication origins and telomeric sequences [40,42]. Accordingly, numerous works suggest that G4 structures can regulate several essential processes in the cell, such as gene transcription, DNA replication, DNA repair, telomere stability and V(D)J recombination [40]. For instance, in mammals, telomeric DNA consists of TTAGGG repeats [39]. They can form G4 structures that inhibit telomerase activity responsible for maintaining length of telomeres and are associated with most cancers [8, 43]. G4s can also regulate gene expression such as for MYC oncogene where inhibition of the activity of NM23-H2 molecules, that bind to the G4, silences gene expression [7]. Moreover, G4s are also fragile sites and prone to DNA double-strand breaks [30]. Accordingly, G4s are highly suspected to be implicated in human diseases such as cancer or neurological/psychiatric disorders [1,11,20].

Following the Human Genome project [13], computational algorithms were developed to predict the location of G4 sequence motifs in the human genome [31,35]. First algorithms consisted in finding all occurrences of the canonical motif *G*_3+_*N*_1–7_*G*_3+_*N*_1–7_*G*_3+_*N*_1–7_*G*_3+_ (or the corresponding C-rich motif) [24, 25]. Using this canonical motif, over 370 thousand G4s were found in the human genome. Nonetheless, such pattern matching algorithms lacked flexibility to accomodate for possible divergences from the canonical pattern. To tackle this issue, novel score-based approaches were developed to compute G4 propensity score by quantifying G-richness and G-skewness [4], or by summing the binding affinities of smaller regions within the G4 and penalizing with the destabilizing effect of loops [22]. A machine learning approach was proposed to predict G4s based on sequence features (such as k-mer occurrences) and trained using newly available genome-wide mapping of G4s in vitro named G4-seq [37]. Deep learning was also recently developed to predict G4s. PENGUINN, a deep convolutional neural network (CNN), was trained to predict G4s in vitro [26]. Another CNN, G4detector, was designed to predict G4s forming both in vitro and in vivo [3]. Thus, all current approaches, except G4detector, aimed to predict G4s in vitro, but were not designed to assess the ability of G4 sequences to form in vivo (*e.g*. G4 activity).

Here, we propose a novel method, named DeepG4, aimed to predict G4 activity from DNA sequence. DeepG4 implements a CNN which is trained using a combination of available in vitro (G4-seq) and in vivo (BG4 ChIP-seq) genome-wide mapping data. DeepG4 achieves excellent accuracy (area under the receiver operating characteristic curve or AUROC > 0.956). DeepG4 outperforms by far all existing non-deep learning algorithms including G4Hunter, pqsfinder and Quadron, and performs better than existing deep learning approaches G4detector and PENGUINN. DeepG4 can predict well across cell lines. Moreover, DeepG4 identifies key DNA motifs that are predictive of active G4s. Among those motifs, we found specific motifs resembling the G4 canonical motif (or parts of G4 canonical motif), but also numerous known transcription factors which could play important roles in G4 activity directly or indirectly. In addition, analysis of publicly available SNPs revealed that SNPs increasing predicted G4 activity were associated with higher gene expression, higher H3K4me3 mark and lower DNA methylation.

## 2 Materials and Methods

### 2.1 BG4 ChIP-seq and G4-seq data

We downloaded BG4 ChIP-seq data for HaCaT, K562 and HEKnp cell lines from Gene Expression Omnibus (GEO) accession numbers GSE76688, GSE99205 and GSE107690 [19,21,28]. Data were mapped to hg19. We downloaded processed qG4 ChIP-seq (similar to BG4 ChIP-seq) peaks mapped to hg19 for breast cancer [20]. We downloaded processed G4-seq peaks mapped to hg19 from GEO accession number GSE63874 [9]. We used G4-seq from the sodium (Na) and potassium (K) conditions.

### 2.2 Active G4 sequences

We defined positive DNA sequences (active G4 sequences) as having the propensity to form both in vitro and in vivo G4s as follows. We only kept BG4 ChIP-seq peaks overlapping with G4-seq peaks. We then used the 201-bp DNA sequences centered on the BG4 ChIP-seq peak summits.

As negative (control) sequences, we used sequences randomly drawn from the human genome with sizes, GC, and repeat contents similar to those of positive DNA sequences using genNullSeqs function from gkmSVM R package (https://cran.r-project.org/web/packages/gkmSVM).

### 2.3 quadparser

We scanned DNA sequences for G4 canonical motifs with pattern *G*_3+_*N*_1–7_*G*_3+_*N*_1–7_*G*_3+_*N*_1–7_*G*_3+_ (or the corresponding C-rich pattern) as implemented in quadparser algorithm [24, 25].

### 2.4 G4CatchAll

We scanned DNA sequences for G4s by considering atypical features using G4CatchAll algorithm [15].

### 2.5 G4Hunter, pqsfinder, qparse and GQRS mapper

We computed G4 propensity score using G4Hunter with default parameters: threshold = 1.5 and width = 25 [4]. We also computed G4 propensity score using pqsfinder, using qparse and using GQRS mapper with default parameters [5, 22, 33]. For a given sequence, several G4 motifs can be detected. In order to obtain only one score per sequence, we used the score sum.

### 2.6 Quadron

We used machine learning approach Quadron to compute G4 propensity [37].

**Figure 1.**
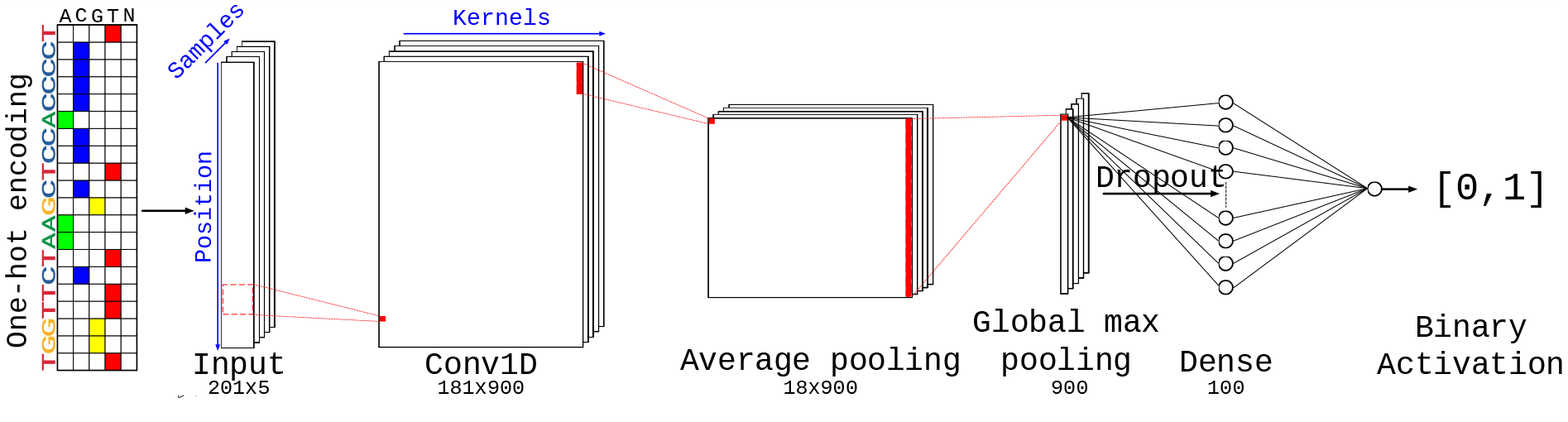
DeepG4 model architecture.

### 2.7 PENGUINN and G4detector

We used deep learning approaches PENGUINN and G4detector to compute G4 score [3,26]. For a fair comparison with DeepG4, PENGUINN and G4detector were retrained using Keras. For each model, the best AUC was picked among the epochs.

### 2.8 JASPAR DNA motifs

We used PWMs for transcription factor binding sites from the JASPAR 2018 database (http://jaspar.genereg.net). We assessed motif enrichment using AME program from MEME suite (http://meme-suite.org/).

### 2.9 SNP data

We downloaded SNPs associated with gene expression from GTEx project (https://www.gtexportal.org/home/datasets). We downloaded SNPs associated with chromatin modifications from Delaneau *et al*. [14]. We also downloaded SNPs associated with DNA methylation from Pancan-meQTL database [18].

### 2.10 DeepG4 model

DeepG4 is a feedforward neural network composed of several layers illustrated in 1. DNA sequence is first encoded as a one-hot encoding layer. Then, a 1-dimension convolutional layer is used with kernels. A local average pooling layer is next used. Then, the global max pooling layer extracts the highest signal from the sequence. Dropout is used for regularization. A dense layer then combines the different kernels and the activation sigmoid layer allows to compute the score between 0 and 1 of a sequence to be an active G4. The model is described in details in Subsection Results and Discussion, Deep learning approach.

Hyperparameters including the number of kernels (900), kernel size (20bp), kernel activation (relu), pool size (10bp), drop-out (0%), epoch number (20), number of neurons in the dense layer (100) and the optimizer choice (rmsprop) were found by fine-tuning with a random grid.

### 2.11 DNA motifs from DeepG4

The first layer of DeepG4 contained kernels capturing specific sequence patterns similar to DNA motifs. For a given kernel, we computed activation values for each positive sequence. If a positive sequence contained activation values above 0, we extracted the sub-sequence having the maximum activation value. The set of sub-sequences was then used to obtain a position frequency matrix (PFM) by computing the frequency of each DNA letter at each position for the kernel.

Each kernel PFM was then trimmed by removing low information content positions at each side of the PFM (threshold > 0.9). PFMs whose size were lower than 5 bases after trimming were removed. PWMs were next computed from PFMs assuming background probability of 0.25 for each DNA letter as done in JASPAR.

Because many PWMs from DeepG4 were redundant, we used the motif clustering program matrix-clustering from RSAT suite (http://rsat.sb-roscoff.fr/) with parameters: median, cor=0.6, ncor=0.6. We used PWM cluster centers as DNA motifs for further analyses.

### 2.12 Predicting SNP effect on G4 activity using DeepG4

We used DeepG4 to predict the effect of a SNP on G4 activity as follows. We computed a Δ*score* as the difference between the DeepG4 score of the sequence with the SNP (alternative allele) and the DeepG4 score of the sequence without the SNP (reference allele). In order to account for the SNP context, G4 activity was predicted from the 201-base sequence surrounding the SNP. Instead of using DeepG4 last layer output (a score varying between 0 and 1) as a score, we used the output from the second to the last layer to obtain a score varying from —∞ to +∞, allowing a better measure for Δ*score*.

### 2.13 DeepG4 implementation and sequence availabity

DeepG4 was implemented using Keras R library (https://keras.rstudio.com/). DeepG4 is available at https://github.com/morphos30/DeepG4. All fasta files used for training and predictions were also deposited.

### 2.14 Comparisons with other G4 tools and availabity

Comparisons with other G4 prediction tools used in this article can be run using a pipeline and a docker available at https://github.com/morphos30/DeepG4ToolsComparison. The docker includes all algorithms, but also retrained algorithms and modified algorithms.

## 3 Results and Discussion

### 3.1 Deep learning approach

Our computational approach, called DeepG4, for predicting active G4 sequences is schematically illustrated in Figure 2. In the first step (Figure 2A), we retrieved recent genome-wide mapping of in vitro G4 human sequences using G4-seq data [9] and of in vivo G4 human sequences using BG4 ChIP-seq data (BG4 is an antibody binding G4s) [19]. Both methods mapped G4s at high resolution (few hundred base pairs). By overlapping BG4 ChIP-seq peaks with G4-seq peaks, we could identify a set of G4 peaks that were formed both in vitro and in vivo, and which we called BG4-G4-seq peaks and considered as “active G4s”.

**Figure 2.**
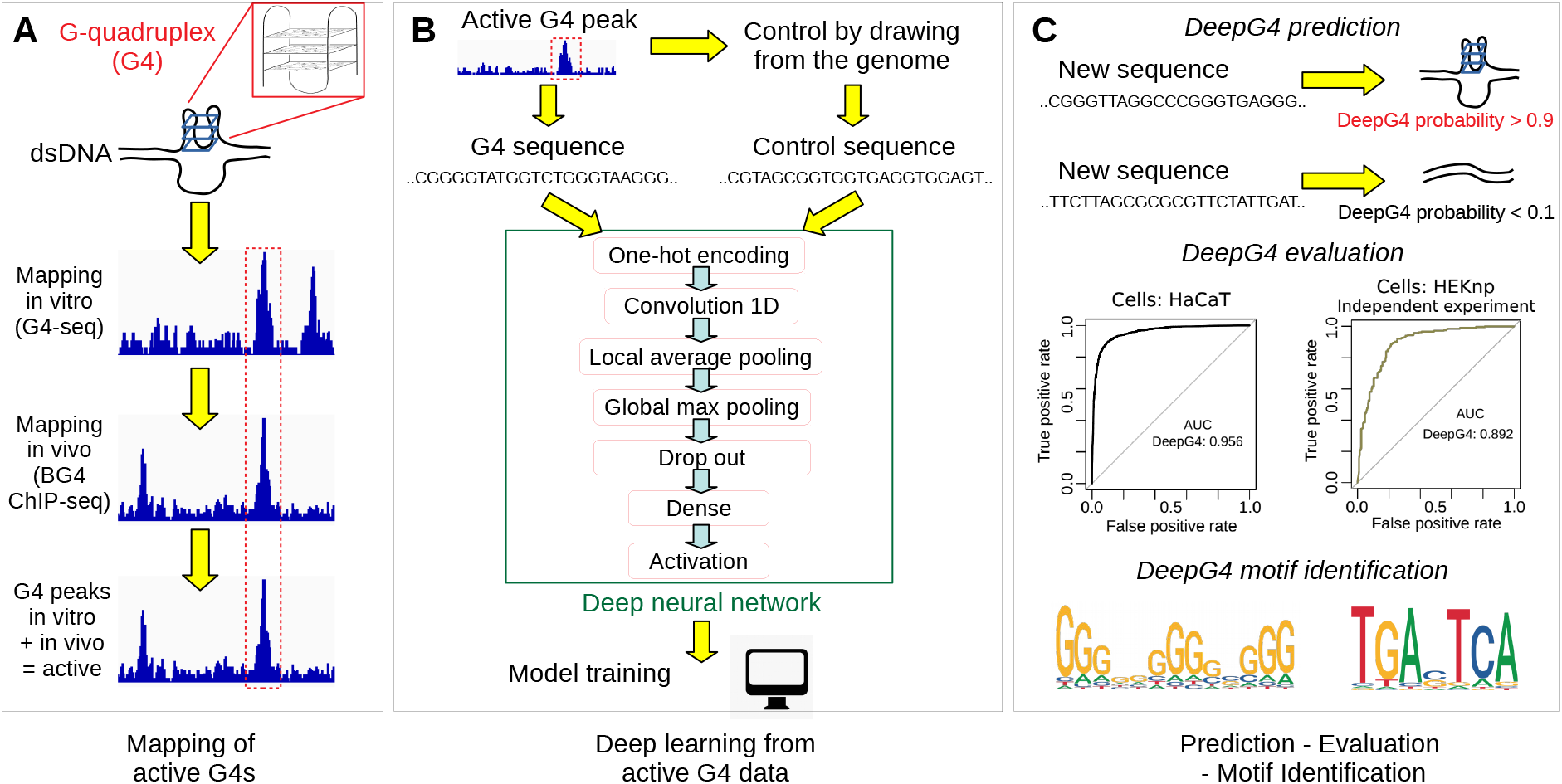
Illustration of DeepG4. A) Mapping of active G4 sequences both in vitro and in vivo using NGS techniques. B) Deep learning model training using active G4 and control sequences. C) G4 activity prediction, evaluation and motif identification.

In the second step (Figure 2B), we extracted the DNA sequences from active G4 peaks (positive sequences). As negative sequences, we used sequences randomly drawn from the human genome with sizes, GC, and repeat contents similar to those of positive DNA sequences. Positive and negative sequences were then used to train our deep learning classifier called DeepG4. DeepG4 is a feedforward neural network composed of several layers. DNA sequence is first encoded as a one-hot encoding layer. Then, a 1-dimension convolutional layer is used with 900 kernels and a kernel size of 20 bp. A local average pooling layer with a pool size of 10 bp is next used. This layer is important: it allows to aggregate kernel signals that are contiguous along the sequence, such that a G4 sequence can be modeled as multiple contiguous small motifs containing stretches of Gs. For instance, a G4 sequence can be defined by two contiguous motifs GGGNNNGGG separated by 5 bases, yielding the canonical motif GGGNNNGGGNNNNNGGGNNNGGG. Then, the global max pooling layer extracts the highest signal from the sequence for each kernel. Dropout is used for regularization. A dense layer then combines the different kernels. The activation sigmoid layer allows to compute the score between 0 and 1 of a sequence to be an active G4.

In the third step (Figure 2C), we used DeepG4 to predict the G4 activity (score between 0 and 1) for a novel DNA sequence. We split the sequence data into a training set to learn model parameters, a validation set to optimize hyper-parameters and a testing set to assess model prediction accuracy. For this purpose, we computed the receiver operating characteristic (ROC) curve and the area under the ROC (AUC). DeepG4 motifs are identified from the convolutional layer.

### 3.2 G4 predictions with DeepG4

Using DeepG4, we obtained excellent predictions of active G4s from HaCaT cells on the test set (Figure 3A; AUC = 0.956; 95% confidence interval of AUC: [0.953, 0.960]). Prediction performance of DeepG4 on an independent dataset with the same cell line (HaCaT) also showed very high accuracy (AUC=0.95, CI95%: [0.946, 0.954]; Figure 3B). We then evaluated the ability of DeepG4 trained on one cell line (HaCaT) to predict G4s in another cell line. We found that DeepG4 could predict reasonably well active G4s in HEKnp cell line [19] (AUC=0.892, CI95%: [0.861, 0.923]; Figure 3C) and in K562 cell line too [29] (AUC=0.92, CI95%: [0.917, 0.924]; Figure 3D).

**Figure 3.**
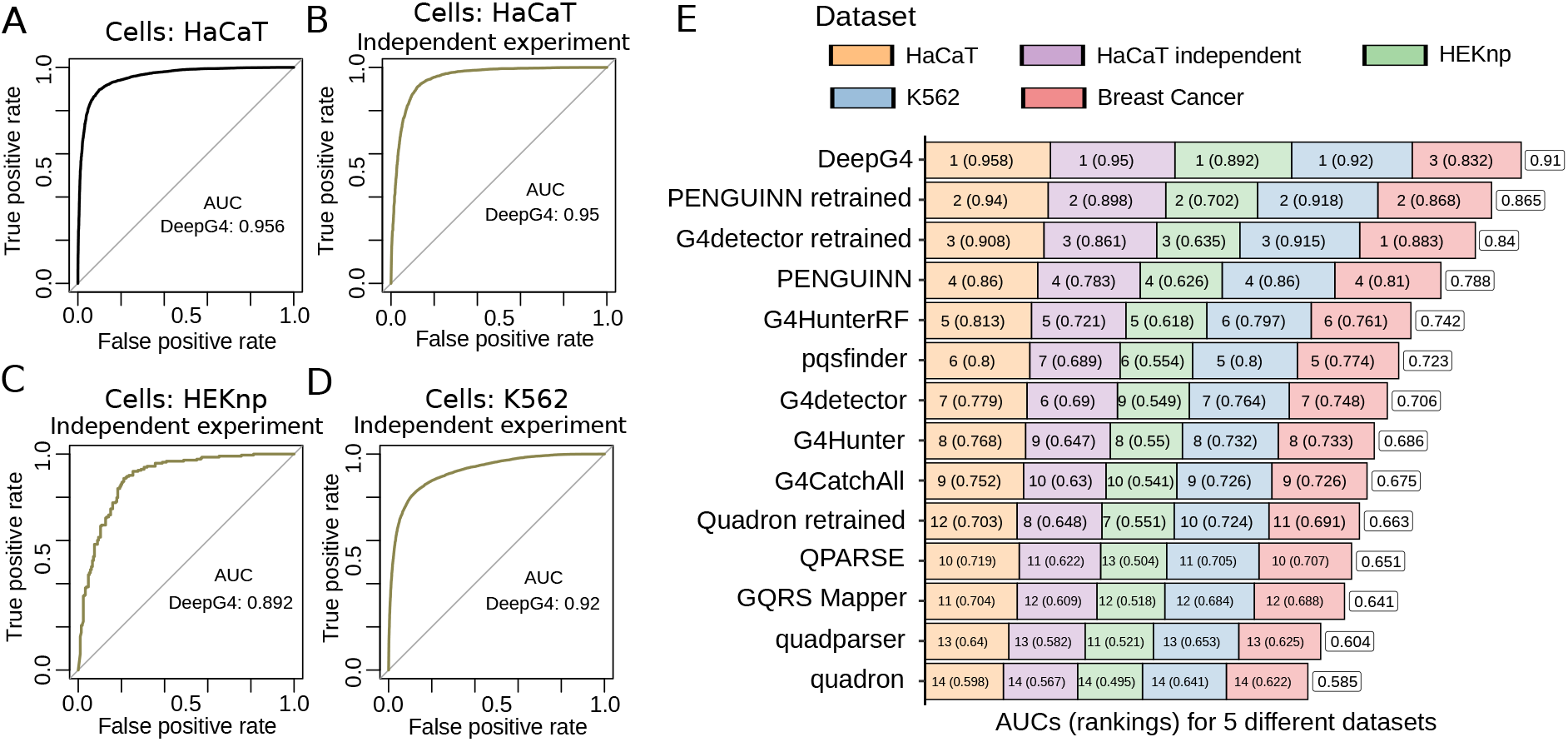
Prediction accuracy of DeepG4 and comparison with state-of-the-art algorithms. A) Prediction accuracy of DeepG4. The model was trained and evaluated using HaCaT cell data (with train and test sets). Receiver operating characteristic (ROC) curve and area under the ROC curve (AUC) were plotted. B) Prediction accurary of DeepG4 trained using one experiment (GEO dataset GSE76688) and evaluated using another independent experiment (GEO dataset GSE99205) in HaCaT cells. C) Prediction accurary of DeepG4 trained using HaCaT data and evaluated in another cell line, HEKnp. D) Prediction accurary of DeepG4 trained using HaCaT data and evaluated in another cell line, K562. E) Comparison of DeepG4 with existing algorithms on 5 different datasets. For each dataset, the AUC ranking for an algorithm is plotted with the AUC value in parentheses. On the right, the average AUC value is plotted for every algorithm.

We then compared DeepG4 with state-of-the-art G4 prediction algorithms, previously shown to perform very well for in vitro G4 predictions. All the datasets, results and algorithms are available from a github repository (see Subsection Comparisons with other G4 tools and availability). DeepG4 outperformed the other algorithms on 4 out of 5 datasets, with an average AUC of 0.91. Deep learning algorithms PENGUINN and G4detector, that we retrained for fairer comparison, ranked 2nd and 3rd with average AUCs of 0.865 and of 0.840, respectively. In particular, they performed the best on the breast cancer data. Among non-retrained algorithms, PENGUINN ranked 4th with an average AUC of 0.788, and G4detector ranked 7th with an average AUC of 0.706. Machine-learning approach Quadron, that was originally trained from in vitro G4s, performed the worst to predict active G4s (average AUC of 0.585, ranked 14th). Re-trained Quadron yielded greatly improved results (average AUC of 0.663, ranked 10th), but which were still far less accurate than those from DeepG4 and other deep learning methods. Lower accuracy of Quadron, which was designed for in vitro G4s, reveals that features aimed to model in vitro G4s were not sufficient to capture the complexity of active G4s.

Score-based algorithms including pqsfinder, G4Hunter, QPARSE and GQRS mapper did not perform well with average AUCs of 0.723 (ranked 6), of 0.686 (ranked 8), of 0.651 (ranked 11), and of 0.641 respectively (ranked 12). Lower performance can be partly attributed to the lack of parameter training of score-based algorithms. To illustrate such drawback of score-based algorithms, we designed a modified version of G4Hunter. G4Hunter depends on one threshold parameter which might not be properly set for active G4s (set to 1.5 as in [4]). Hence, we implemented G4Hunter within a machine learning approach, such that G4Hunter threshold parameter could be learned from data, thereby allowing a fairer comparison with DeepG4. For this purpose, we ran G4Hunter predictions for varying G4Hunter threshold values (1 to 2 by 0.1 increment) as features for a random forest classifier (we named G4HunterRF). G4HunterRF strongly outperformed previous G4Hunter predictions with an average AUC of 0.742, but the predictions were still less good than those from DeepG4 and deep learning methods in general. Motif-based algorithms, such as quadparser and G4CatchAll, also performed poorly when predicting active G4s with average AUCs of 0.604 and of 0.675.

These results thus demonstrated the strong ability of DeepG4 to accurately predict active G4s from DNA sequences, and the better performances compared to retrained existing deep learning approaches. Moreover, results also showed the incapacity of state-of-the-art non-deep learning algorithms to predict precisely G4s that were formed in vivo. In addition, we showed that DeepG4 could accurately predict G4s in other cell lines.

### 3.3 Identification of important motifs from DeepG4

The first layer of DeepG4 convolutional neural network comprised kernels that encoded DNA motifs predictive of active G4s. Hence, we extracted from the first layer the kernels and converted them to DNA motif PWMs to better understand which motifs were the best predictors of G4 activity. DeepG4 identified 900 motifs, many of them were redundant. To remove redundancy, we clustered the motifs using RSAT matrix-clustering program and kept the cluster motifs (also called root motifs in the program) for subsequent analyses. Cluster motifs could be divided into two groups: a group of de novo motifs and a group of motifs that resembled known TFBS motifs. To distinguish between these two groups, we used TomTom program (MEME suite) which mapped the cluster motifs to JASPAR database. DeepG4 motifs matching JASPAR were considered as known TFBS motifs, while motifs that did not match were classified as de novo motifs.

We then sought to assess the ability of DeepG4 motifs to predict G4s. Hence, we computed DeepG4 cluster motif variable importances using random forests and found strong predictors (Figure 4A). In order to visualize the cluster motifs on a map, we used multi-dimensional scaling (MDS), where we also added the original kernel motifs used to build the cluster motifs. We found that the first MDS component reflected the motif GC content (higher at the left side), while the second component represented the guanine stretch length (higher at the top) (Figure 4B).

**Figure 4.**
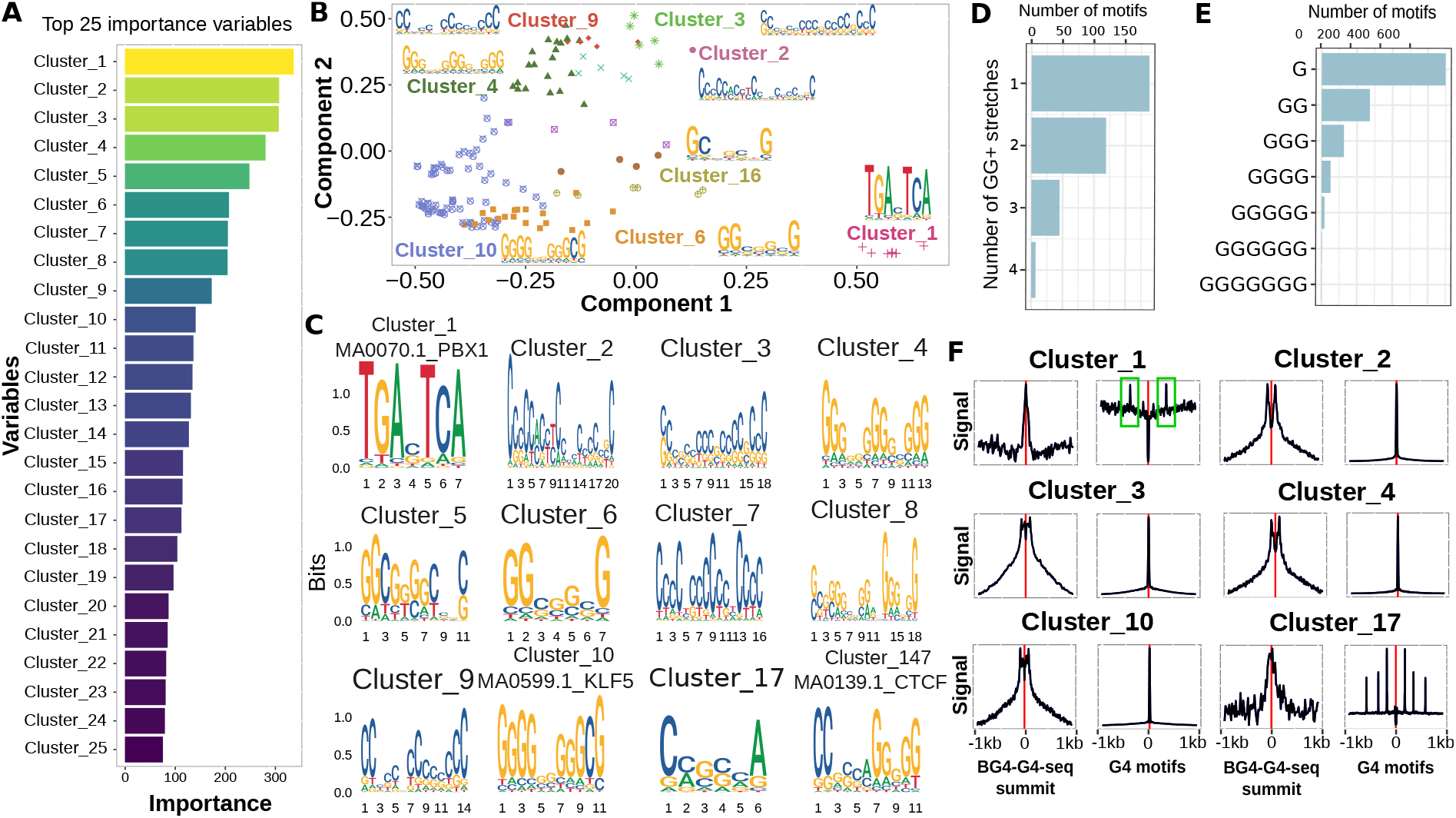
DNA motifs identified by DeepG4. A) Variable importances of DeepG4 cluster motifs, as estimated by random forests. Clustering of DeepG4 kernel motifs was done by RSAT matrix-clustering program to obtain cluster motifs. B) Multidimensional scaling (MDS) of DeepG4 motifs. As an input, matrix-clustering correlation matrix between kernel motifs was used. C) Logos of cluster motifs with highest variable importances. D) Number of kernel motifs containing one or more GG+ stretches. A GG+ stretch is defined as a stretch of 2 or more Gs in the motif consensus sequence. E) Number of kernel motifs containing G stretches depending on stretch length. F) Average profiles measuring the enrichment of cluster motifs centered around BG4-G4-seq summits or canonical G4 motifs.

Many strong predictors were de novo motifs which ressembled parts of G4 canonical motifs. For instance, cluster 4 comprised 3 stretches of GGG+, and could thus be considered as 3/4 of a canonical G4 motif (Figure 4C). Cluster 5 comprised two stretches of GG+, thus forming half of a canonical G4 motif. We then counted GG+ stretches (stretches of 2 or more guanines) from the kernel motifs and found that many kernel motifs contained more than one GG+ stretch (Figure 4D). Moreover, the guanine stretches were of varying lengths, ranging from one G up to 5 Gs (Figure 4E).

Among the best predictors, we also found several motifs corresponding to known TFBS motifs (Figure 4C). For instance, the best predictor, cluster 1, almost perfectly matched PBX1 motif MA0070.1 (q-value=0.011). Another strong predictor, cluster 10, matched KLF5 motif MA0599.1 (q-value=0.016). We could also identify a predictor, cluster 147, that matched well CTCF motif (q-value=0.0001), but whose variable importance was moderate.

We then assessed the enrichment of DeepG4 cluster motifs around BG4-G4-seq summits and around canonical G4 motifs. Motifs ressembling parts of G4 canonical motifs, such as clusters 2, 3 and 4, were enriched at both BG4-G4-seq summits and canonical G4 motif centers, thus representing actual G4 structures. But other motifs that were very different from the G4 canonical motif, such as cluster 1, were strongly enriched at BG4-G4-seq summits, but depleted at the exact location of canonical G4 motifs. Interestingly, cluster 1 was enriched close to canonical G4 motifs (around 300 bp, highlighted in green), suggesting that cluster 1 (PBX1 motif MA0070.1) did not participate directly to the G4 structure, but could act in the vicinity to support G4 activity. Similarly, cluster 17 motif, which did not resemble G4 parts nor matched a TF, showed enrichment at BG4-G4-seq summits, but was enriched in the vicinity of canonical G4 motifs. Moreover, we could consider a third class of cluster motifs, such as cluster 10, which matched a TF (KLF5) and was found strongly enriched at BG4-G4-seq summits and at the exact location of canonical G4 motifs.

These observations revealed the important role of TFBS motifs that could act directly in G4 activity as part of G4 structure, as previously shown for SP1 in vitro [36], or could participate indirectly to support G4 activity in the vicinity of G4s as for PBX1 motif.

### 3.4 TF motif predictors of active G4s vary between cell types

We then wanted to explore if TFBS motifs that are predictors of active G4s may vary between cell types. For this purpose, we built random forest classifiers to distinguish between active G4s from one cell type and active G4s from the other cell types (for instance HaCaT active G4s against other active G4s), using all known TFBS motifs as predictors. The random forest for HaCaT active G4s obtained an AUC of 0.885. Similarly, AUCs of 0.604 and of 0.711 were obtained for K562 and breast cancer specific active G4s. Thus, the TFBS motifs were able to partly predict cell type specific active G4s.

We then estimated variable importances from the random forests. The motifs corresponding to the top-10 highest importances for each cell line were shown in Figure 5A (only the motifs enriched in the cell line active G4s were shown). We could identify many cell type specific predictors. For instance, motif MA0476.1, corresponding to FOS, was the best motif to discriminate HaCaT (keratinocyte) active G4s from other active G4s (Figure 5B). FOS is a member of the AP-1 complex whose TFs are well-known key regulators of epidermal keratinocyte survival and differentiation [16]. In K562 (myelocytic leukemia) cells, EWSR1-FLI1 motif MA0149.1 was the best predictor (Figure 5B). Among the best predictors, we also identified SPI1 (motif MA0080.5) known to activate gene expression in myeloid and B-lymphoid cell development [32]. In breast cancer cells, ZNF354C motif MA0130.1 was the best discriminator (Figure 5B), respectively. The third best predictor was ETV4 (motif MA0764.2), which was involved in triplenegative breast cancer [45].

**Figure 5.**
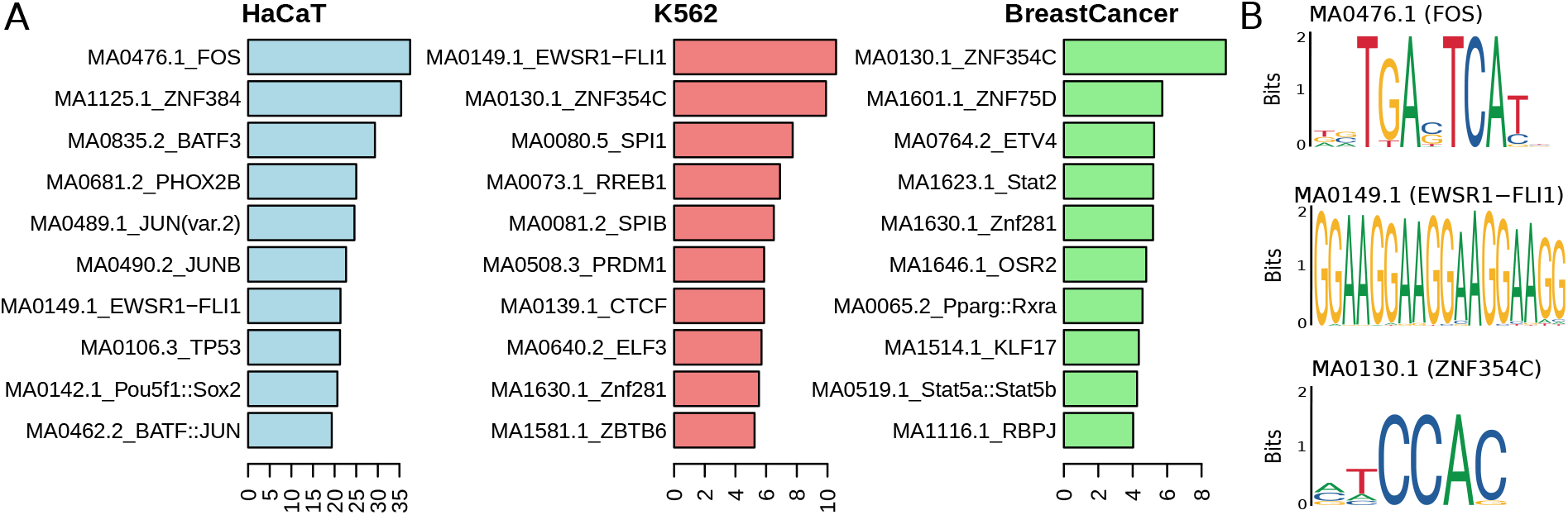
Cell type specific transcription factor motif predictors of active G4s. A) Variable importances of transcription factor (TF) motifs were computed using random forests to distinguish between active G4s from one cell type and active G4s from the other cell types. B) For each cell line, the motif logo of the first TF motif predictor of active G4s.

Results thus support the role of TFs as key regulators of G4 activity, depending on cell type. In particular, TFs could influence the activity of G4s during cell differentiation and proliferation.

### 3.5 SNP effect on g-quadruplex

We then sought to explore the possible effect of SNPs on G4 activity using DeepG4, as a previous study showed that SNPs within G4 canonical motifs could affect gene expression [2]. For this purpose, we predicted G4 activity from the 201-base sequence surrounding a SNP. To assess the effect of a SNP, we computed a delta score as the difference between the DeepG4 score of the sequence with the SNP (alternative allele) and the DeepG4 score of the sequence without the SNP (reference allele).

We first analyzed the link between delta score and gene expression, and found that SNPs increasing gene expression presented higher G4 activity as compared to SNPs diminishing gene expression (*p* = 9 × 10^-15^; Figure 6A). We then explored the link between G4 activity and chromatin mark H3K4me3 which was often associated with the activation of transcription of nearby genes. We found that SNPs increasing H3K4me3 mark were associated with higher delta score as compared to SNP diminishing gene expression (*p* = 3 × 10^-5^; Figure 6B). Next, we looked at DNA methylation which is often involved in the inhibition of nearby genes. Moreover, G4 structures are known to affect DNA methylation [29]. In agreement, we observed that SNPs reducing DNA methylation showed higher G4 activity compared to SNPs increasing DNA methylation (p = 3 × 10^-5^; Figure 6C).

**Figure 6.**
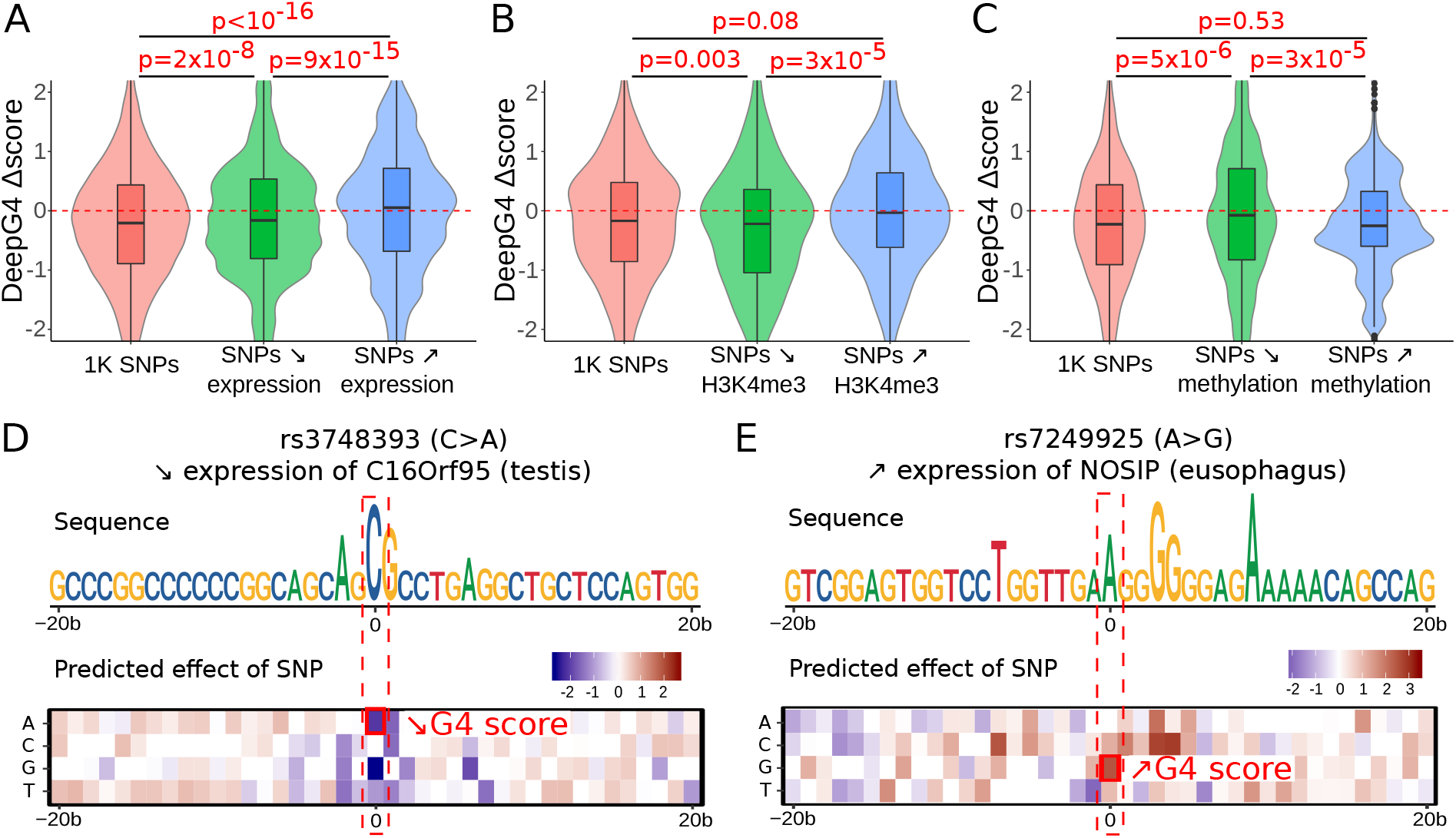
G-quadruplex SNPs are associated with gene expression and chromatin activity. A) SNP effect of g-quadruplex (delta score) depending on SNP effect on gene expression (regression beta). SNPs are from GTEx database [12]. SNPs from 1000 genomes project (1K SNPs) were added as controls. B) SNP effect of g-quadruplex (delta score) depending on SNP effect on H3K4me3 mark (regression beta). SNPs are from Delaneau *et al*. [14]. C) SNP effect of g-quadruplex (delta score) depending on SNP effect on DNA methylation (regression beta). SNPs are from Pancan-meQTL database [18]. D) Example of the SNP rs3748393 that is associated with gene expression decrease and which is predicted to decrease G4 activity. The predicted effect of SNPs for each allele and each position around rs3748393 is plotted as heatmap. E) Example of the SNP rs7249925 that is associated with gene expression increase and which is predicted to increase G4 activity.

We next illustrated SNP effects on G4s with 2 examples. The SNP rs3748393 had a C allele as reference and A allele as alternative, and was associated with C16Orf95 gene repression in testis (gene expression regression 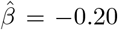). Using DeepG4, we found that alternative allele A led to a decrease in G4 activity (Figure 6D). In fact, the SNP was part of a G4 motif composed of several G stretches (C stretches in the figure). Thus, by changing the C allele to A (corresponding to G>T on the reverse strand), the SNP could alter the G4 structure stability. As a second example, we then analyzed SNP rs7249925 that was associated with NOSIP gene induction in eusophagus (gene expression regression 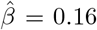). We found that the alternative allele G, which expanded a GGGGGG stretch, led to an increase of G4 activity (Figure 6E).

These results suggest that SNPs do not only affect gene expression, DNA methylation and transcription factor binding, but also affect secondary structures of DNA such as G4s and that affect in turn many molecular processes including transcription and chromatin dynamics.

## 4 Conclusion

In this article, we propose a novel deep learning method, named DeepG4, to predict active G4s from DNA sequences. The proposed method is designed to predict G4s that are found both in vitro and in vivo, unlike previous algorithms that were developed to predict G4s forming in vitro (naked DNA). DeepG4 outperforms all existing non-deep learning algorithms even when those algorithms were (re-)trained using active G4 data. This is due to the DeepG4 convolutional layer that allows to learn predictive features (here DNA motifs) from the DNA sequence without prior knowledge. Compared to current deep learning algorithms, DeepG4 also performs better even after retraining the model.

Moreover, DeepG4 uncovered numerous specific DNA motifs predictive of active G4s. Many motifs resembled the canonical G4 motif (*G*_3+_*N*_1–7_*G*_3+_*N*_1–7_*G*_3+_*N*_1–7_*G*_3+_) or even parts of it. Most notably, many motifs corresponded to half or 3/4 of the canonical motif. The combination of these G4 parts, which is captured by DeepG4, brings flexibility in G4 modeling. Strikingly, many other motifs matched known TFBS motifs including PBX1 motif MA0070.1, KLF5 motif MA0599.1 and CTCF motif MA0139.1, suggesting that they could contribute directly to G4 structures themselves or participate indirectly in G4 activity in the vicinity through the binding of transcription factors. In agreement, it was previously found that G4s are enriched in the vicinity of the architectural protein CTCF at 3D domain (topologically associating domain, TAD) borders [23]. Moreover, it has been shown that SP1 binds to G4s with a comparable affinity as its canonical motif [36].

Analysis of SNP effects on predicted G4s suggested that SNPs altering G4 activity could affect transcription and chromatin, *e.g*. gene expression, H3K4me3 mark and DNA methylation. In particular, we showed that SNPs decreasing G4 activity tended to inhibit gene expression, in agreement with many results supporting a positive link between G4s and transcription [41]. Interestingly, our results also revealed that SNPs diminishing G4 activity also induced H3K4me3 decrease. Conversely, SNP altering G4 activity increased DNA methylation, which is also supported by recent results showing the tight link between G4 structure and hypomethylation at CpG islands in the human genome [28]. In the context of genome-wide association studies (GWASs), DeepG4 thus paved the way for future studies assessing the impact of known disease-associated variants on DNA secondary structure and provides a novel mechanistic interpretation of SNP impact on transcription and chromatin.

There are several limitations of the proposed approach. First, the prediction accuracy of DeepG4 strongly depends on existing datasets that are limited, especially regarding in vivo mapping. Once more techniques for in vivo G4 mapping will be developped, DeepG4 will need to be retrained in order to improve prediction accuracy. Moreover, since DeepG4 was trained based on human data, predictions on non-mammalian genomes are expected to be inaccurate. Second, DeepG4 is limited to predict active G4s but a similar approach could be used to predict any active non-B DNA structure using permanganate/S1 nuclease footprinting data [27]. Third, DeepG4 only uses DNA sequence as input. Incorporating chromatin accessibility with DNA-seq (or ATAC-seq) as input to DeepG4 could allow the model predict cell-type specific active G4s.

## Authors’ contributions

VR conceived and implemented the model, analyzed the SNPs and wrote the Github reposity. MG analyzed DNA motifs from DeepG4 and carried out complementary SNP analysis. EN generated control sequences and compared DeepG4 with existing algorithms. RM designed the project, conceived the model and wrote the manuscript.

## Acknowledgements

The authors are grateful to Balasubramanian lab for data (University of Cambridge, UK). The authors are also thankful to Matthias Zytnicki for comments.

